# rEXPAR: an isothermal amplification scheme that is robust to autocatalytic parasites

**DOI:** 10.1101/518118

**Authors:** Georg Urtel, Jean-Christophe Galas, André Estevez-Torres

## Abstract

In the absence of DNA, a solution containing the four deoxynucleotidetriphosphates (dNTPs), a DNA polymerase and a nicking enzyme generates a self-replicating mixture of DNA species called parasite. Parasites are problematic in template-based isothermal amplification schemes such as EXPAR, as well as in related molecular programming languages, such as the PEN DNA toolbox. Here we show that the nicking enzyme Nb.BssSI allows to change the sequence design of EXPAR templates in a way that prevents the formation of parasites when dATP is removed from the solution. This method allows to make the EXPAR reaction robust to parasite contamination, a common feature in the laboratory, while keeping it compatible with PEN programs, which we demonstrate by engineering a parasite-proof bistable reaction network.

---

Isothermal nucleic acid amplification methods (1) are interesting alternatives to the polymerase chain reaction (PCR) because they do not need thermocycling equipment and are thus suited for the detection of nucleic acids in resource-limited environments (2). Different molecular implementations exist, such as nucleic acid sequence-based amplification (NASBA), strand displacement amplification (SDA), rolling circle amplification (RCA), loop-mediated isothermal amplification (LAMP) and exponential amplification reaction (EXPAR) (3).

Among the cited methods EXPAR has two important advantages. Firstly, it has a short detection time in the order of minutes (4). Secondly, it produces single stranded DNA (ssDNA) as an output which can be used, either to set up simple colorimetric detection methods based on the aggregation of DNA-decorated nanoparticles (5), or to couple EXPAR to molecular programs capable of displaying complex spatiotemporal dynamics (6, 7, 8, 9). EXPAR exponentially amplifies the concentration of a trigger ssDNA **A** in the presence of a ssDNA template **T**, a DNA polymerase and a nicking endonuclease, called nickase in the following (Figure 1a).

**Figure 1.**
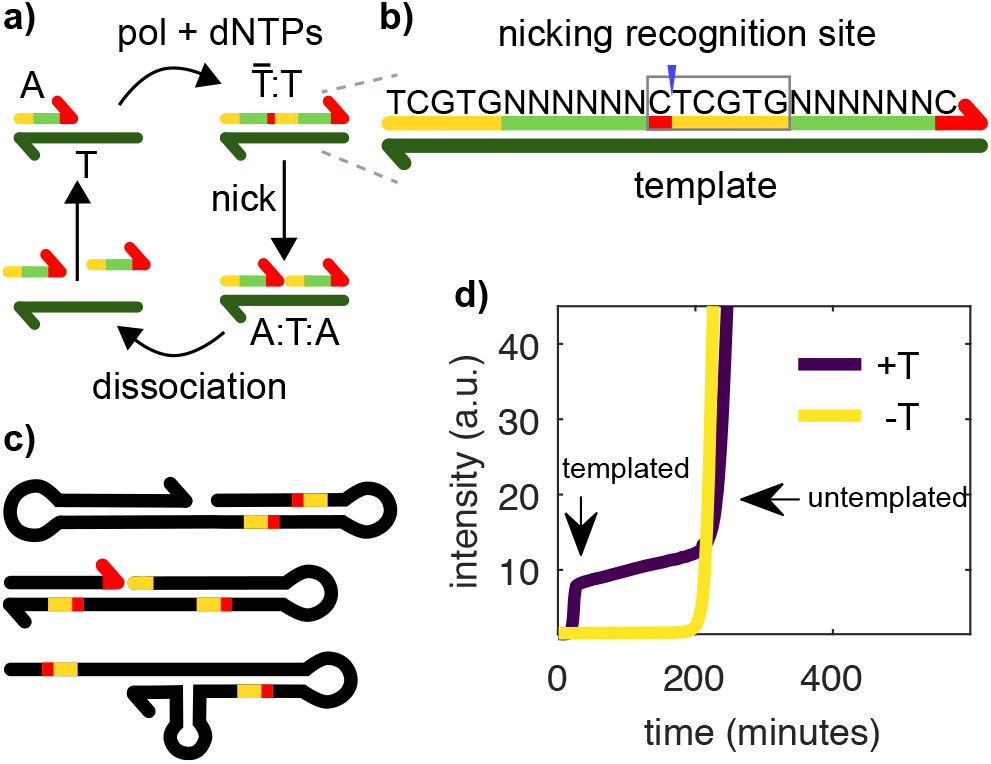
Templated and untemplated replication in the EXPAR reaction. a) During templated replication, the trigger **A** is elongated on the template **T** by a polymerase (pol), consuming dNTPs. The double stranded complex 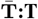 is nicked by a nickase (nick) and two **A**s are created, which can dissociate and replicate on other **T**s. b) Each trigger contains a split-up recognition site for the nickase (red and yellow), which is completed after elongation. The sequence of the six N nucleotides can be chosen freely. c) In EXPAR experiments an untemplated replicator, termed the parasite, emerges after some time. The parasite is not a single sequence, but a pool of sequences. They are rich in secondary structures and bear recognition sites for the nickase. Unlike shown here, they can reach several kbp in length. d) EvaGreen fluorescence *vs*. time for an EXPAR reaction in the presence (**+T**, purple) and in the absence (**– T**, yellow) of template strand **T**_1_.

However, EXPAR has two important drawbacks induced by non-specific reactions. The first one, usually known as early phase background amplification (4), or self-start, limits the detection of very low quantities of DNA. Several solutions have recently been proposed for this problem (10, 11, 12). The second problem, known as late-phase background amplification (4), or untemplated amplification, arises in systems where nucleic acids are exponentially amplified with the help of enzymes. Mutations lead to new sequences and, after some time, a parasitic sequence — or set of sequences—emerges, which is able to replicate more efficiently than the initial target sequence. One famous example is Sol Spiegelman’s monster, that arouse during the *in vitro* replication of Q*β*-RNA with Q*β*-replicase and nucleotides (13). The EXPAR reaction produces a different kind of parasitic species containing repetitive and palindromic sequences where, typically, AT tracts are flanked by nickase recognition sites (4). Importantly, parasites appear by *de novo*, or untemplated, synthesis of DNA. Autocatalytic parasites have been observed in the presence of enzymes other than nickases, such as restriction enzymes (14), helicases (15) and possibly T7 RNA polymerase (16), and also in PCR reactions either as a side-effect (17) or by design (18).

Because parasite replication is as efficient as templated replication (SI Figure S4), parasites easily contaminate the whole laboratory and are difficult to eradicate, thus posing a problem to make EXPAR a robust analytical technique. Furthermore, when EXPAR is used to build more complex molecular programs, such as oscillators or bistable networks, the emergence of parasites limits the lifetime of these systems to typically one day, restricting the use of these powerful molecular programs (19) for building nonequilibrium materials (9). In this work, we show that choosing a particular nickase, Nb.BssSI, allows to change the sequence design of EXPAR templates in a way that prevents the formation of parasites when dATP is removed from the solution. We further demonstrate that this approach permits the detection of DNA in the presence of contaminating parasites and that it is compatible with the design of bistable reaction networks.

## METHODS

Oligonucleotides were purchased from IDT and their sequences displayed in Tables 2 and S1. Template strands were HPLC purified and trigger strands were desalted. The enzymes we used were 8-40 U/mL Bst DNA Polymerase, Large Fragment (NEB), 20-500 U/ml Nb.BssSI (NEB) and 0 or 100 nM of in-house produced *thermus thermophilus* RecJ exonuclease (20). We noticed a 3.4-fold batch-to-batch change in Nb.BssSI activity. The reaction buffer contained 20 mM Tris-HCl, 10 mM (NH_4_)_2_SO_4_, 50 mM NaCl, 1 mM KCl, 2 mM MgSO4, 6 mM MgCl_2_, 1 g/L synperonic F 108 (Sigma-Aldrich), 4 mM dithiothreitol, 0.1 g/L BSA (NEB), 1x EvaGreen Dye (Biotium) and 0.1x ROX (invitrogen). Nucleotides (NEB or invitrogen) were added in different concentrations and compositions. In some experiments netropsin was added as indicated. Experiments were performed in a reaction volume of 20 *μ*L at 44 °C, with 50 nM template concentration, when necessary, in a CFX96 Touch Real-Time PCR Detection System (Bio-Rad) or a Qiagen Rotor-Gene qPCR machine. The intensity of the green channel was recorded every minute. To avoid cross-contaminations, experiments involving pipetting of solutions containing parasites were performed in a different room using a different set of pipets.

## RESULTS AND DISCUSSION

### Suppressing untemplated replication

In EXPAR templated amplification the trigger species **A** replicates on template **T** with the help of a polymerase and a nickase, noted pol and nick respectively in the following (Figure 1a). There is a series of sequence constrains for the reaction to happen. If we note *a* the sequence of **A**, and *ā* its complementary, **T** is a double repeat of *ā*, noted *āā*. In addition, the double stranded species 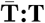 bears the recognition site of the nickase in such a way that the enzyme cuts 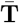 to generate two species **A** bound to **T** (Figure 1b). As a result, when **A** binds to the 3’ end of **T** it is extended by pol to make 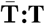, which is cut by nick to form the complex **A:T:A**. The reaction is isothermal and set up close to the melting temperature of **A:T** to ensure that A can dehybridize and take part in reactions with other templates. To close the catalytic loop, either **A:T:A** dissociates to recycle species **T** or it is extended again by pol, which is capable of strand-displacement, regenerating 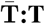 and producing an extra **A**.

Untemplated replication in the presence of pol, nick and dNTPs is well documented (4, 21) but its mechanism has not yet been elucidated (22). In the current working hypothesis, as soon as a sequence with a hairpin on the 3’ appears by *de novo* synthesis, it may be extended by pol. Subsequent rounds of hairpin formation and slippage followed by polymerization account for the synthesis of palindromic repetitive sequences. Nicking events followed by the novo polymerization of few bases that can self-hybridize may explain the observation of palindromic sequences flanked by nicking sites (Figure 1c).

Figure 1d displays a typical EXPAR experiment where the total concentration of double stranded DNA (dsDNA) is followed by recording the fluorescence of a dsDNA intercalator. A solution containing pol, nick, dNTPs and template strand **T**_1_ was incubated at constant temperature (purple line). In the beginning of the reaction, the signal coming from the dye is low since no dsDNA is present. When templated replication occurs, at 18 min, a rapid exponential phase leads to a signal increase until the signal saturates, indicating that all template is bound to the trigger. After 200 minutes, a second, non-linear signal increase takes place, which is due to untemplated replication. In the absence of template (yellow line), only the later signal increase is observed. Interestingly, the onset time of untemplated replication is similar in the presence or in the absence of template.

A strategy used so far to mitigate parasite replication was the addition of netropsin, since the parasite sequence was found to be rich in AT repeats by Tan et al. (4, 19). Netropsin helps by binding such AT-rich sequences, but cannot always prevent parasite formation, probably because other studies showed parasites without AT-rich stretches (21). Our solution to the problem consists in making it impossible to create secondary structures that can be nicked. Table 1 shows the sequences of the recognition sites for commercially-available nickases. Most of the enzymes, including Nt.BstNBI a common nickase used in EXPAR, contain all four bases (A, C, G, T) in their recognition site, and thus both templated and untemplated replication need the four dNTPs to proceed. In contrast, Nb.BssSI has a recognition site with only three bases (C, G, T). As a result, one can design a trigger with only C, G, T and a template with only C, G, A in their sequences. In the absence of dATP, such a system should be able to perform templated replication normally, while being uncapable of untemplated replication. Indeed, if mutations lead to the formation of unwanted products, they might contain secondary structures, but no recognition site for the nickase, which needs two adenines. This unwanted sequence will then only be able to grow by extending the 3’-end, which is not an exponential process and should’t interfere too much with autocatalysis.

**Table 1.**
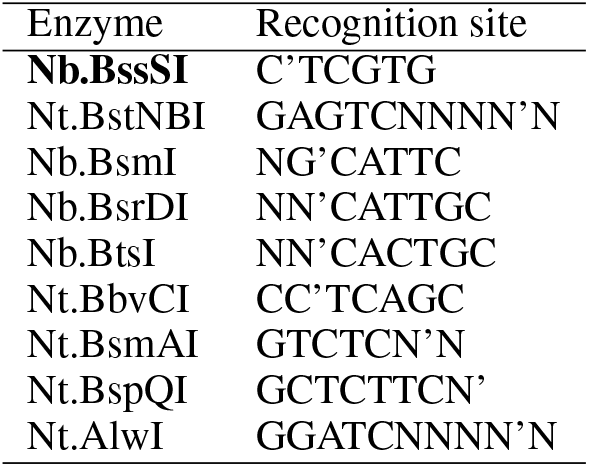
Recognition sequences of different nicking enzymes (from 5’ to 3’). The nicking sites are marked with’. N means any base. In bold the enzyme used in this study.

Figure 2 demonstrates that this approach works as designed. We incubated template **T**_1_ with pol and nick in the presence or in the absence of dATP. Because we will later use this system to design more complex molecular programs, template **T**_1_ lacked two bases in the 3’ side compared to the complementary of a double repeat of the trigger sequence **A**_1_, i.e. **A**_1_ was a 12-mer, **T**_1_ a 22-mer and when **A**_1_ binds to **T**_1_ and is extended by the polymerase the 24-mer **A**_1_**A**_1_ is formed (Table 2). In addition, ttRecJ, a single-stranded specific exonuclease with 5’ activity, noted exo, was added to the reaction to ensure that template replication was active throughout the duration of the experiment. **T**_1_ was protected from the exonuclease by 3 phosphorothioate (PTO) bonds at its 5’ end and bear a fourth one in the middle of the sequence to protect against nick star activity (see below). In these conditions, when dATP was present, both templated and untemplated replication were observed, while only templated amplification was observed in the absence of dATP (Figure 2a). Analysis of the reaction products on a denaturating polyacrylamide gel confirmed that at long times, large amounts of strands much longer than the trigger, which we identify with the parasite, only form when dATP is present (Figure 2b). Before the emergence of the parasite, the reaction products were indistinguishable both in the presence or in the absence of dATP: 3 bands at 12, 22 and 24-nt corresponding respectively to **A**_1_, **T**_1_ and **A**_1_ **A**_1_. After 200 min —when the parasite has already appeared in the presence of dATP— no new bands appear in the –dATP sample. Finally, it is straightforward to design trigger and template sequences compatible with parasite suppression. Figure 2c shows three sets of sequences (SI Table S1) that displayed templated but not untemplated replication in the absence of dATP. This last set of experiments was performed in the absence of exo, to demonstrate that degradation is not needed for parasite suppression. Because this approach allows to perform EXPAR reactions that are robust against untemplated replication we will call it rEXPAR in the following.

**Figure 2.**
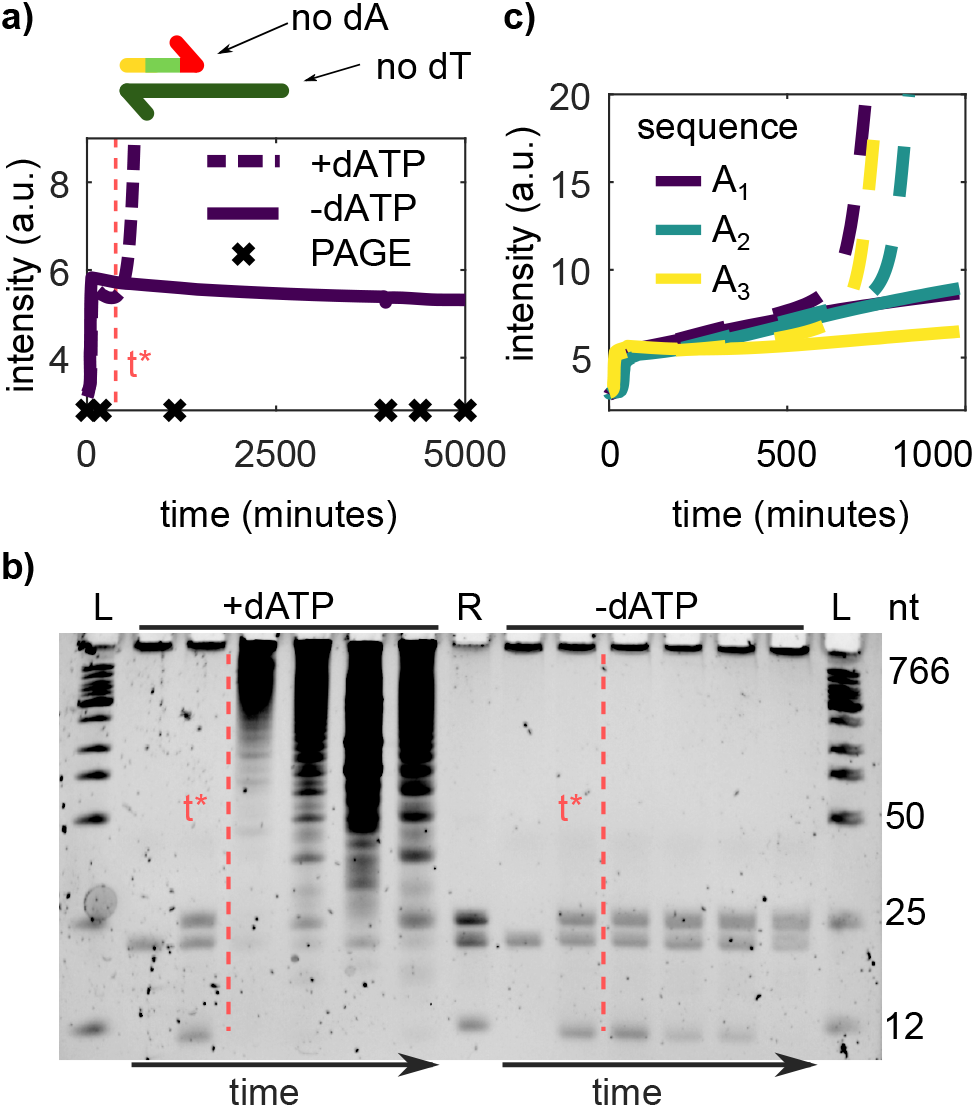
Suppressing dATP blocks untemplated replication without perturbing templated replication. a) Top: We use a nickase that allows to design templates without thymine (dT) and triggers without adenine (dA). Bottom: EvaGreen fluorescence vs. time for an EXPAR reaction of trigger **A**_1_ and template **T**_1_ in the presence (dashed) and in the absence (solid line) of dATP. Crosses indicate time-points where aliquots were withdrawn for the gel in panel (b). b) Time evolution of the reactions in panel (a) on a PAGE denaturing gel. The red dashed line is a guide to the eye indicating the onset of untemplated replication in the presence of dATP (t*). Lanes 4-7 have been diluted 20-fold for easier visualization of parasite bands. L is a ladder and R a reference containing species **A**_1_, **T**_1_ and **A**_1_**A**_1_. c) EXPAR reactions for three different sequences in the presence (dashed) and in the absence (solid line) of dATP. Conditions: 8 (a,b) or 4.8 (c) U/mL pol, 20 U/mL nick, 100 (a,b) or 0 (c) nM exo, 1 (a,b) or 0.4 (c) mM dNTPs.

**Table 2.**
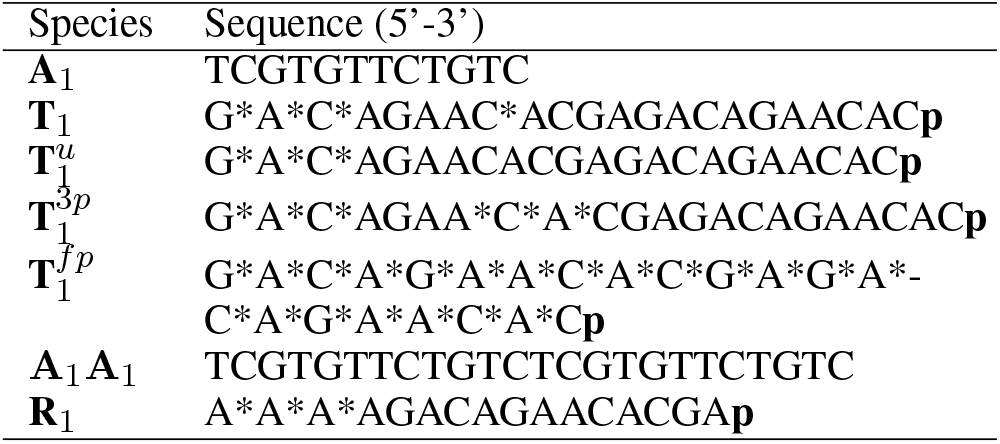
DNA sequences for the main set of species used in this work. Asterisks indicate phosphorohioate bonds and p phosphate modifications.

### Nb.BssSI nickase displays a star activity that can be easily suppressed

Nb.BssSI, the nickase used here has been recently developed and has seldom been used in EXPAR experiments to our knowledge. In our preliminary experiments we used template 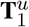, identical to **T**_1_ except for a missing PTO after the 8th nucleotide, and we observed a significant loss of fluorescence signal in many long-term experiments (Figure 3b purple line). In EXPAR experiments with other nicking enzymes (Nb.BsmI, Nt.BstNBI) we did not observe this behavior. With 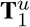, increasing Nb.BssSI concentrations promoted long-term signal loss while the addition of 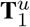 temporarily restored the signal (SI Figure S1). We hypothesized that, because Nb.BssSI was derived from the restriction enzyme BssSI, it may have a reminiscent restriction activity, which would cleave the template strand. To solve this problem, we tested different templates with additional PTO protection (note that all 5’-ends have 3 PTOs for protection from exonuclease). Figure 3b shows EXPAR experiments with the protected templates. The unprotected template 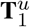 shows degradation after only 20 min. The fully protected template 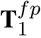 does not only have the PTOs in the recognition sequence, but between all 22 bases. This leads to a strong inhibition of templated replication that could arise from the higher melting temperature of PTO-modified strands or from the inhibition of pol or nick. In contrast, templates with 1, **T**_1_, or 3, 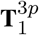, PTOs surrounding the putative second nicking site associated to the BssSI activity on the template strand show almost uninhibited replication and a stable signal in the steady state for at least 5000 minutes (Figure 2a).

**Figure 3.**
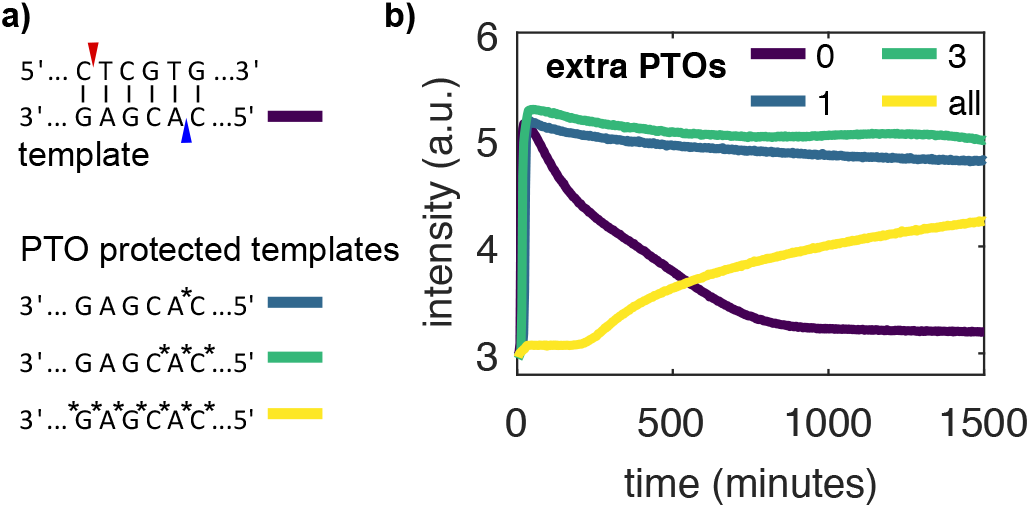
Nb.BssSI star activity can be suppressed by adding a PTO bond to the template strand. a) The nickase Nb.BssSI is supposed to cut at the site indicated in red. The restriction enzyme BssSI additionally cuts at the site indicated in blue. We tested four templates, 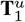 (purple), **T**_1_ (blue), 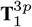 (green), 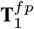 (yellow) with increasing number of PTOs around the BssSI site. b) EvaGreen fluorescence *vs*. time for rEXPAR experiments with templates in (a). Conditions: 8 U/mL pol, 200 U/mL nick, 100 nM exo, 0.4 mM dNTPs.

### Standard methods delay parasite emergence but do not suppress it

Standard methods to mitigate untemplated replication involve decreasing pol or increasing nick concentrations (9) and adding netropsin (4, 19). Figure 4 shows that these approaches delay but do not suppress the onset of parasite emergence. In the absence of template, the parasite onset time, *τ_u_*, is inversely proportional to pol concentration, as expected from first-order kinetics (Figure 4a,b). In contrast, increasing nick concentration increases *τ_u_*, specially at low pol concentration (SI Figure S2), suggesting that nick inhibits pol. Finally, adding up to 4 *μ*M netropsin delays parasite emergence 4-fold but it also slows down templated replication by the same amount and reduces the fluorescence signal from the dsDNA intercalator dye (Figure 4c and SI Figure S3). In summary, all these methods slow down untemplated replication at the cost of slowing down templated replication as well, which is an undesirable feature for rapid analysis.

**Figure 4.**
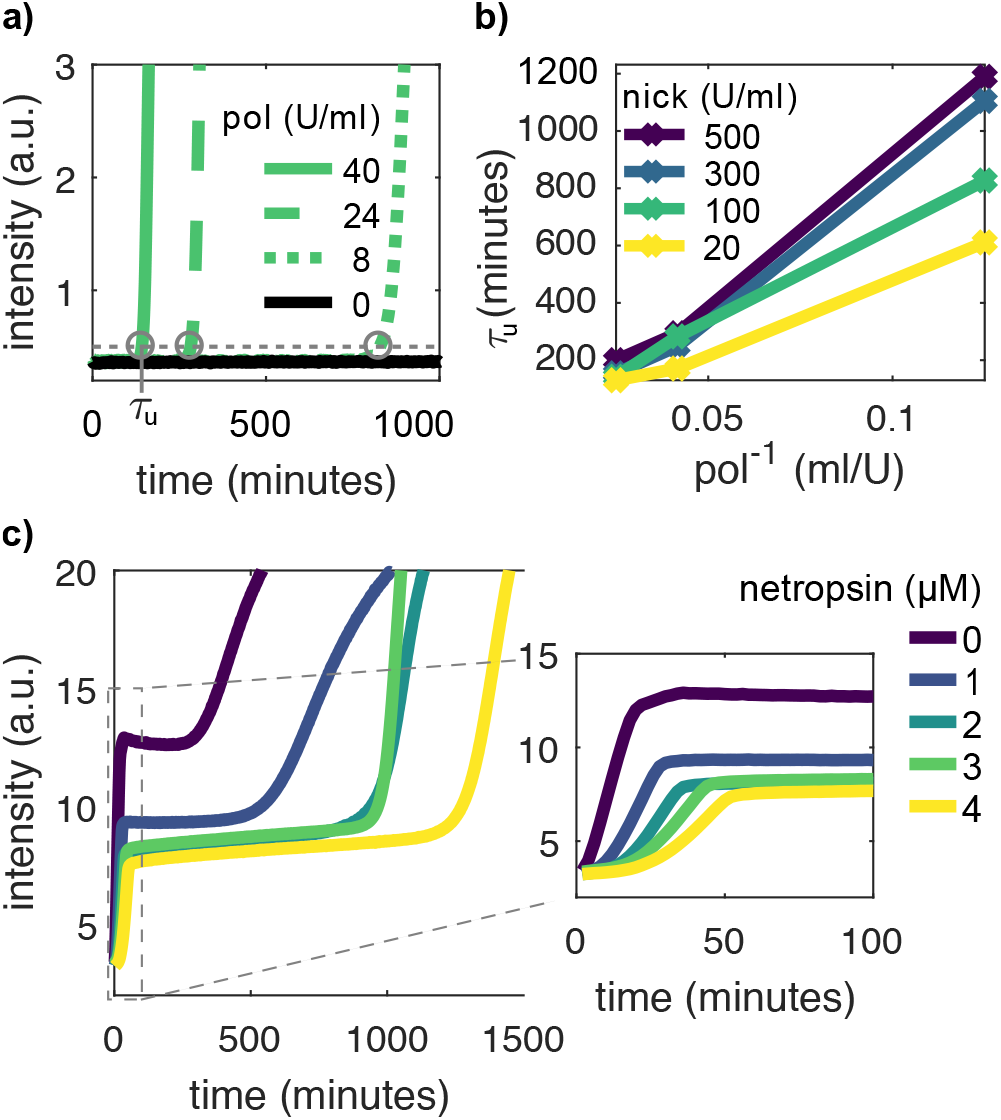
Standard approaches delay untemplated replication but do not suppress it. a) EvaGreen fluorescence vs. time in the absence of **T**_1_ for different polymerase (pol) concentrations at 100 U/ml nickase. Circles indicate the onset time of untemplated replication *τ_u_*. b) *τ_u_ vs*. the inverse of pol concentration for different nick concentrations. c) EvaGreen fluorescence vs. time in the presence of **T**_1_ for different netropsin concentrations. Conditions: 0.4 mM dNTPs, 100 nM exo (for a-b) and 8 U/mL pol, 60 U/mL nick (for c).

### rEXPAR is compatible with PEN molecular programs

An EXPAR autocatalytic network is an intrinsically monostable dynamical system. This means that, in the absence of trigger strand (*i.e. A = 0*), an infinitesimally small addition of **A** will grow exponentially. This is problematic to detect very low amounts of **A**, as any unprimed synthesis of **A** will result in an undesired background, known as early stage background amplification or self-start. One can make EXPAR robust to self-start by using instead a bistable autocatalytic network based on the polymerase, nickase, exonuclease dynamic network assembly toolbox (PEN DNA toolbox) (10).

The PEN DNA toolbox is an experimental framework based on the EXPAR reaction that allows designing reaction networks that mimick the dynamics of gene regulatory networks in solution (6, 19, 23). This framework makes network design straightforward because network topology is defined by predictable interactions between short ssDNAs. In addition, the combination of its three core enzymes with large quantities of dNTPs provides a convenient way to keep the network out of equilibrium in a closed reactor for very long time at steady state. These two unmatched properties have allowed the rational design of complex spatio-temporal behaviors such as oscillations (6, 24), bistability (7, 10) and reaction-diffusion patterns (8, 25, 26, 27). Besides suppressing background amplification in EXPAR, such dynamic behaviors have important applications such as nucleic acid detection (28), material science (9), microrobots (29) and protein directed evolution (30). However, untemplated amplification is an important obstacle for these applications because it precludes the use of PEN circuits for long periods of time.

PEN networks are usually built around one or more autocatalytic nodes based on EXPAR. An exonuclease enzyme is added such that nodes are not only dynamically produced but also degraded. In addition to the autocatalytic template strands intrinsic to EXPAR, that catalyze the reaction **A** → 2**A**, other templates can be used to catalyze other processes such as activation (**A** → **A**+**B**), and repression (**A** →ø). All these template strands bear PTOs in 5’ to protect them from exonuclease degradation.

In this framework, to render bistable an EXPAR reaction, and thus suppress self-start, one just needs to add a second template strand **R** that binds to trigger **A** and, with the help of pol, it extends it into the waste strand **W**, that can be degraded by exo but not recognized by nick (Figure 5a). In addition, the polymerization reaction of **A** on **R** needs to be faster than the one of **A** on its template **T**. To fulfill these two requirements, we chose to use a repressor strand R1 with a sequence of 4 adenines followed by the reverse complementary sequence of **A**_1_ (Table 2) and a template strand that lacks on the 3’ end two bases to be fully complementary to **A**_1_, here 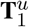. **R**_1_ was protected against exonuclease degradation by 3 PTO bonds in the 5’ end. We incubated 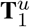 in the presence of pol, nick and exo and increasing concentrations of *R*_1_, noted *R*_1_, in the absence of trigger **A**_1_ and dATP. At the lowest *R*_1_ tested of 15 nM, untriggered templated amplification was observed within 30 min, indicating that the system is monostable with a single stable point at high **A**_1_ concentration. Increasing *R*_1_ resulted in a dramatic increase of the templated amplification time *τ_t_* until it became undetectable (> 1000 min, SI Figure S5) above *R*_1_ = 60 nM (Fig. 5b-c). Above this threshold the reaction network becomes bistable with a stable point at *A*_1_ = 0 and a second one at high *A*_1_. A plot of 1/*τ_t_* as a function of the concentration of **R**_1_ indicates that *R*_1_ is a bifurcation parameter of the reaction network. Because dATP was absent, no untemplated amplification was observed in the system, demonstrating that rEXPAR is compatible with the construction of DNA-based reaction networks with complex dynamics, such as bistability.

**Figure 5.**
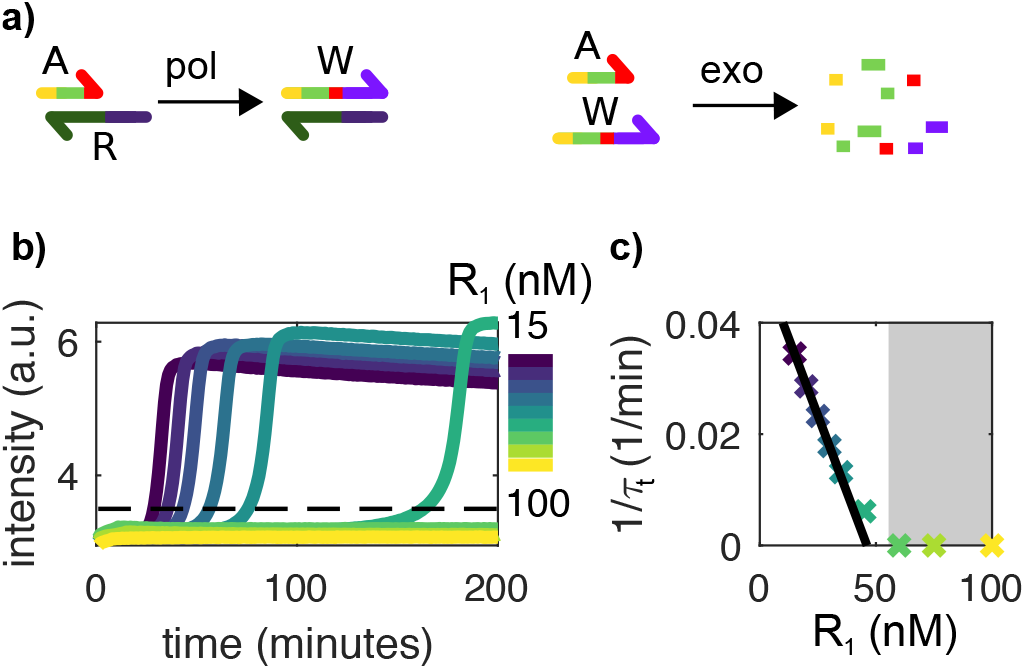
rEXPAR is compatible with PEN DNA bistable programs. a) In addition to the autocatalytic network depicted in Fig. 1a, a repressor template **R** is added that binds to trigger **A** and turns it into waste **W**, which can be degraded by exonuclease but cannot prime autocatalysis. b) EvaGreen fluorescence *vs*. time in the presence of 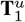 but in the absence of **A**_1_, for increasing concentrations of **R**_1_. The dashed line indicates the threshold corresponding to the onset time of templated amplification *τ_t_*. c) 1 /*τ_t_ vs*. **R** concentration from panel (b). The black line corresponds to a linear fit of slope 1.1 × 10^−3^ nM^−1^min^−1^. The gray area indicates where the system is bistable and thus robust to self-start. Experiments performed in the absence of dATP. Conditions: 8 U/mL pol, 20 u/mL nick, 100 nM exo and 0.4 mM dNTPs.

### rEXPAR is robust to parasite contamination

Besides being an intriguing instance of molecular evolution, parasites can easily contaminate the laboratory (31) and produce false positives in EXPAR because they amplify as fast as target DNA and produce a higher fluorescent signal. Figure 6 shows EXPAR amplification experiments in the presence and in the absence of dATP (rEXPAR) for samples with or without parasite contamination. Contamination consisted in a 3000-fold dilution of another sample that had previously undergone untemplated amplification. This amount corresponds to using a pipet tip that has been filled and emptied with 1 *μ*L of a solution containing parasite (SI Figure S4). We observed that templated and untemplated amplification occurred concomitantly in the sample containing both dATP and parasite. In contrast, rEXPAR samples without dATP were robust to untemplated amplification both in the absence and in the presence of contamination.

**Figure 6.**
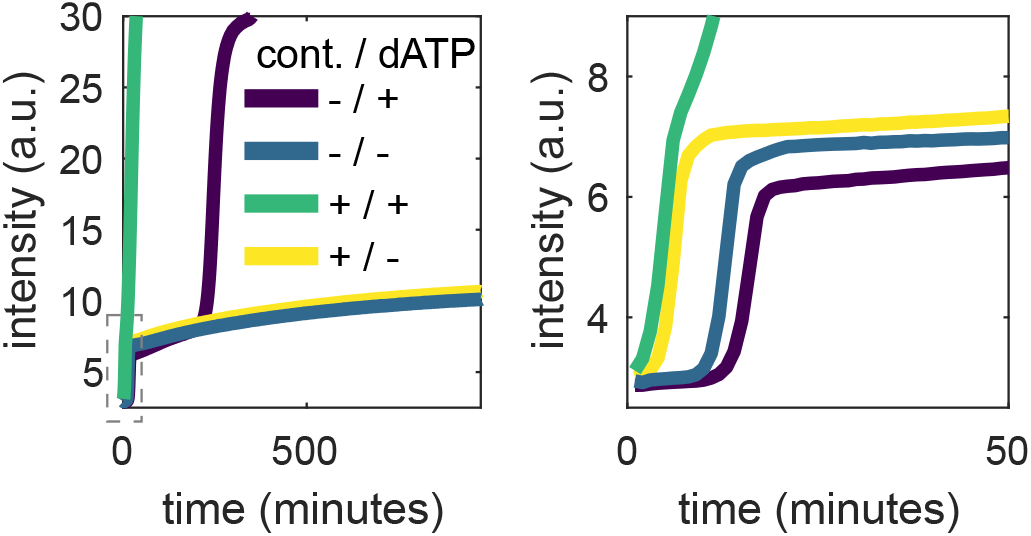
rEXPAR is robust to parasite contamination. EvaGreen fluorescence *vs*. time for EXPAR experiments performed both with and without dATP and contamination from a parasite solution diluted 3000-fold (cont.). Conditions: 40 U/mL po, 500 U/mL nick, 0 nM exo, 60 nM **R**_1_, 0.4 mM dNTPs.

In a second set of experiments (Figure 7 and SI Figure S8) we evaluated the performance of EXPAR and rEXPAR to detect trigger DNA in the presence of parasite contamination. To be more realistic about a fortuitous parasite contamination we used a higher dilution of 10^6^-fold. EXPAR and rEXPAR attained a similar limit of detection of 0.4 pM in the absence of contamination, although rEXPAR was 1.6-fold faster (SI Figures S6-7). In contrast, in the presence of contamination rEXPAR was able to detect the trigger without noticeable change while EXPAR was unable to detect even the highest trigger concentration of 100 pM.

**Figure 7.**
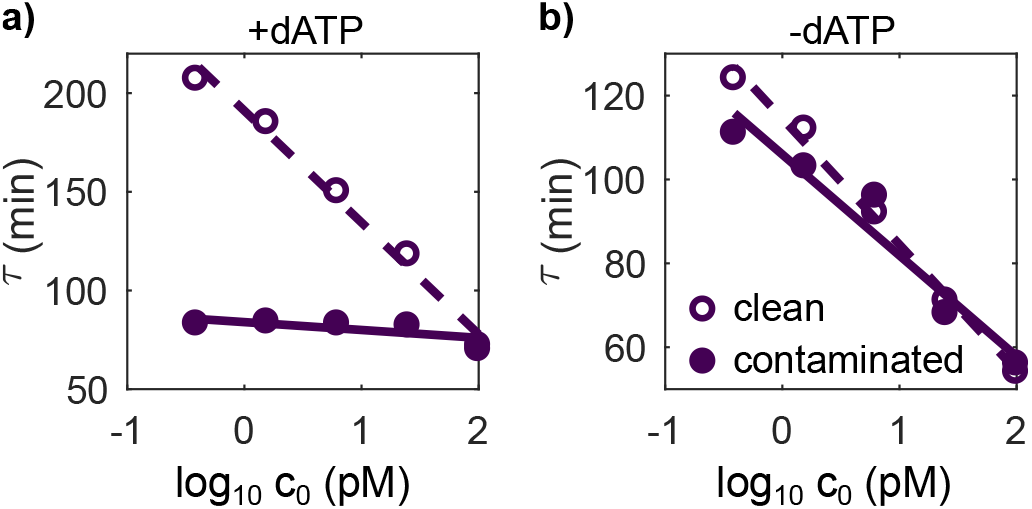
rEXPAR is able to detect trigger DNA in the presence of parasite contamination, in contrast with EXPAR. Amplification onset time, *τ, vs*. the decimal logarithm of the initial trigger concentration co for EXPAR reactions performed in presence (a) and in the absence (b) of dATP, with (filled symbols) and without (empty symbols) contamination from a parasite solution diluted 10^6^-fold. Conditions: 8 U/mL pol, 20 U/mL nick, 100 nM exo, 15 nM **R**_1_, 0.4mM dNTPs.

## CONCLUSION

We took advantage of a new nicking enzyme, Nb.BssSI, which recognition site has only a three-letter code (C’TCGTG) to perform EXPAR isothermal amplification experiments in the absence of one deoxynucleotide (dATP). In these conditions, templated amplification proceeded normally and even slightly faster, while untemplated amplification, resulting in the production of an autocatalytic set of parasitic sequences, was completely suppressed. Our approach, called rEXPAR for robust EXPAR, contrasts with existing methods, such as the addition of netropsin, that mitigate but do not suppress untemplated amplification. rEXPAR is compatible with EXPAR-based molecular programming languages such as the PEN DNA toolbox, which we demonstrated by implementing a bistable autocatalytic network that suppressed self-start spurious reactions. In addition rEXPAR, in contrast with EXPAR, allows the detection of trigger DNA in the presence of minute amounts of parasite contamination. As a result, we believe that rEXPAR will be useful, both for running out-of-equilibrium molecular programs over extended periods of time, which is essential for building “life-like” materials (9), and for making EXPAR more robust in analytical applications.

## SUPPLEMENTARY DATA

The supplementary data contains extra DNA sequences, characterization of the star activity of Nb.BssSI, quantification of netropsin effect on kinetics, parasite replication kinetics, effect of dNTPs concentration on kinetics and amplification curves corresponding to Fig. 7.

## ACKNOWLEDGEMENTS

Yannick Rondelez and Guillaume Ginés for insightful discussions.

