## Supplementary materials for "rEXPAR: an isothermal amplification scheme that is robust to autocatalytic parasites"

### Supplementary information for: rEXPAR: an isothermal amplification scheme that is robust to autocatalytic parasites

### Contents


**1 Sequences** **3**

**2 Nb.BssSI degrades unprotected templates** **3**

**3 Parasite emergence depends on enzyme concentrations** **4**

**4 Netropsin slows down templated replication** **5**

**5 Parasite replication kinetics** **6**

**6 Bistability** **8**

**7 Increasing nucleotide concentration slows down templated amplification** **9**

**8 Amplification curves for EXPAR and rEXPAR in the presence of parasite contamination** **11**

### 1 Sequences

Additional oligonucleotide sequences that do not appear in Table 2 in the MT can be found in Table S1. DNA was purchased from Integrated DNA Technologies, Inc.

Table S1: DNA sequences with modifications. PTOs are marked with an asterisk * and phosphates with **p**.

| **Name** | **Sequences 5' → 3'** |
|---|---|
| $T_1^{+2}$ | G*A*C*AGAACACGAGACAGAACACGA**p** |
| $A_2$ | TCGTGGTTCTTC |
| $T_2$ | G*A*A*GAACCACGAGAAGAACCAC**p** |
| $A_3$ | TCGTGTCTGTC |
| $T_3$ | G*A*C*AGACACGAGACAGACACG**p** |

### 2 Nb.BssSI degrades unprotected templates

During preliminary experiments, we observed a loss of fluorescence signal in many of the long-term experiments, for example in Figure S1a. Two possible reasons for this signal loss are template degradation by Nb.BssSI or a degradation of Nb.BssSI itself. Since the signal loss is stronger for higher concentrations of nickase, a template degradation seemed more likely. To test this assumption, we injected either template or Nb.BssSI to the samples which showed the strongest degradation (Fig. S1b). In addition, the storage medium of template (water) or Nb.BssSI (diluent B, NEB) was injected with the same volumes as a control to compensate for dilution effects. Only the injection of template restores the signal, which starts to slowly decrease again.

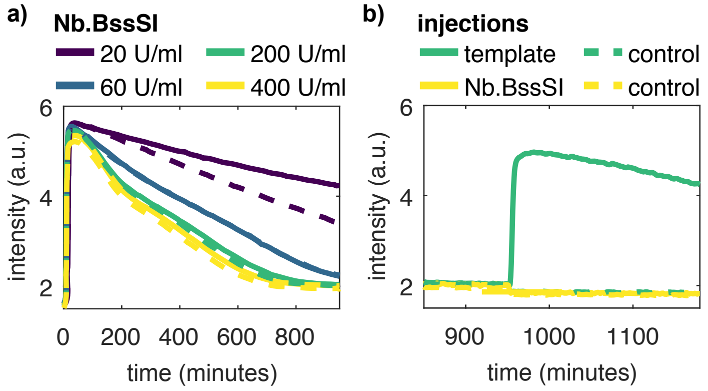


Figure S1: Degradation of templates by Nb.BssSI. a) Autocatalysis with a range of nicking enzyme shows a signal loss over time. Two samples were used for each nickase concentration (solid, dashed). b) For the samples with the strongest signal loss in panel (a), injections were performed. Only injecting template restores the signal. Nickase injections or controls with the respective buffer show no effect.

### 3 Parasite emergence depends on enzyme concentrations

As described in the main text, the emergence of parasites depends on the concentrations of nickase and polymerase as well as the presence of all four dNTPs. Figure S2 show the data represented in Figure 4a-b of the main text plotted in different ways to highlight the effect of nick concentration.

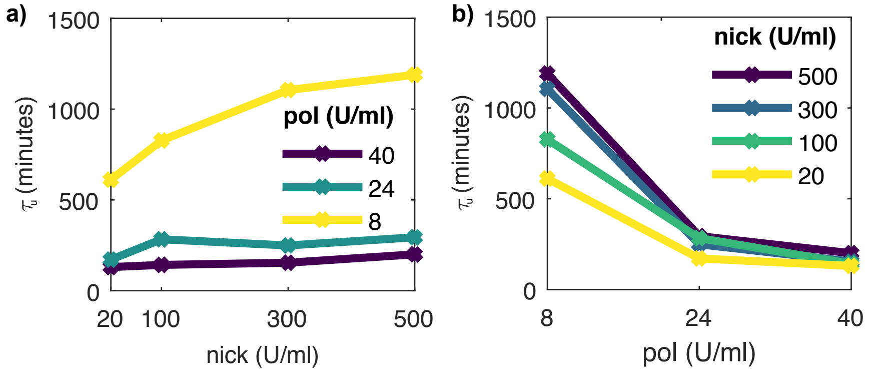


Figure S2: Onset time of untemplated replication $\tau_u$ for different polymerase and nickase concentrations.

### 4 Netropsin slows down templated replication

In Figure 4c of the main text it was shown that netropsin helps to delay the emergence of parasite. Netropsin also has some side effects. It decreases the fluorescence intensity of EvaGreen (Fig. S3a) and slows down the reaction rate (Fig. S3b-d). We found that the rate is proportional to 1/[netropsin]. Comparing the rates for the sample with 4 $\mu$M netropsin to the control sample without netropsin, shows a 4.5-fold decrease in templated replication rate.

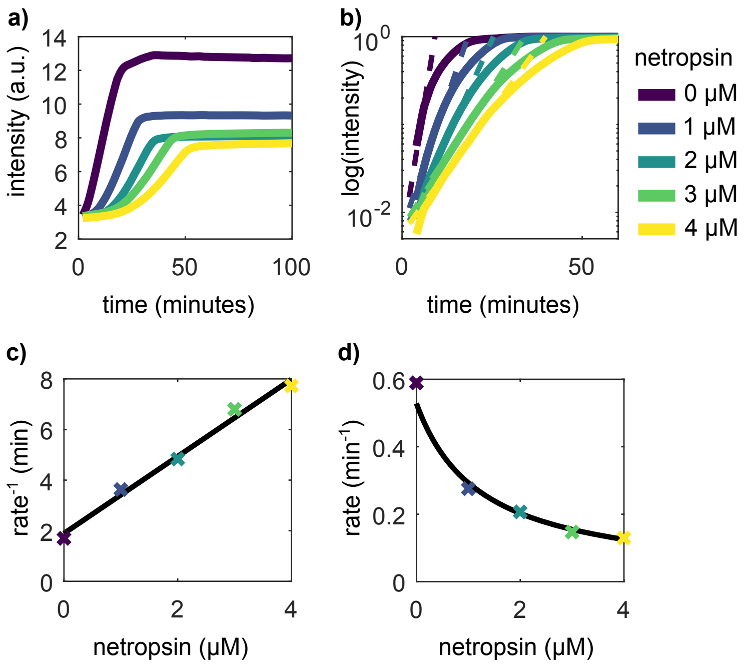


Figure S3: Netropsin inhibits templated replication. The legend for all four plots is on the right side. a) This is the inset of Figure 4c of the main text. It shows that netropsin leads to decreased signal. b) Replotting the data on a log-scale after normalization and fitting exponentials (dashed curves) allows to determine rates. c) Plot of the inversed rates vs. the netropsin concentration and linear fit. The fit shows that the rates decrease with 1/[netropsin]. d) Replot of c) with inversed y-axis.

### 5 Parasite replication kinetics

To study the parasite replication kinetics, we contaminated the reaction mixtures. For the contaminations we used the parasite that emerged in the untemplated reaction shown in Figure 1d of the main text. Figure S4a shows different dilutions of parasite in a reaction mixture with all nucleotides in the absence of template. For each dilution, the onset reaction time $\tau$ was determined and plotted *vs.* the decimal logarithm of the dilution factor. A linear fit to this data provided the untemplated replication rate $r = 0.05\ \mathrm{min}^{-1}$. The enzyme concentrations were the same as for the experiment of Figure 7 of the main text (8 U/ml polymerase, 20 U/ml nickase and 50 nM exonuclease). The rate of templated replication in Figure 7 for the two non-contaminated samples were r=0.04 $min^{-1}$ and r=0.08 $min^{-1}$, respectively in the presence and in the absence of dATP. Thus, the replication rates of the parasite and of trigger $\mathbf{A}_1$ in the presence of 50 nM $\mathbf{T}_1$ are comparable. However, $\mathbf{A}_1$ is a 12-mer, while the parasite is much longer and thus gives much more signal in the real-time cyclers. In Figure S4b we contaminated an untriggered EXPAR experiment with different dilutions of parasite. In the absence of dATP, high concentrations of parasite have an effect on templated self-start. In the presence of dATP, templated self-start and untemplated replication cannot be distinguished.

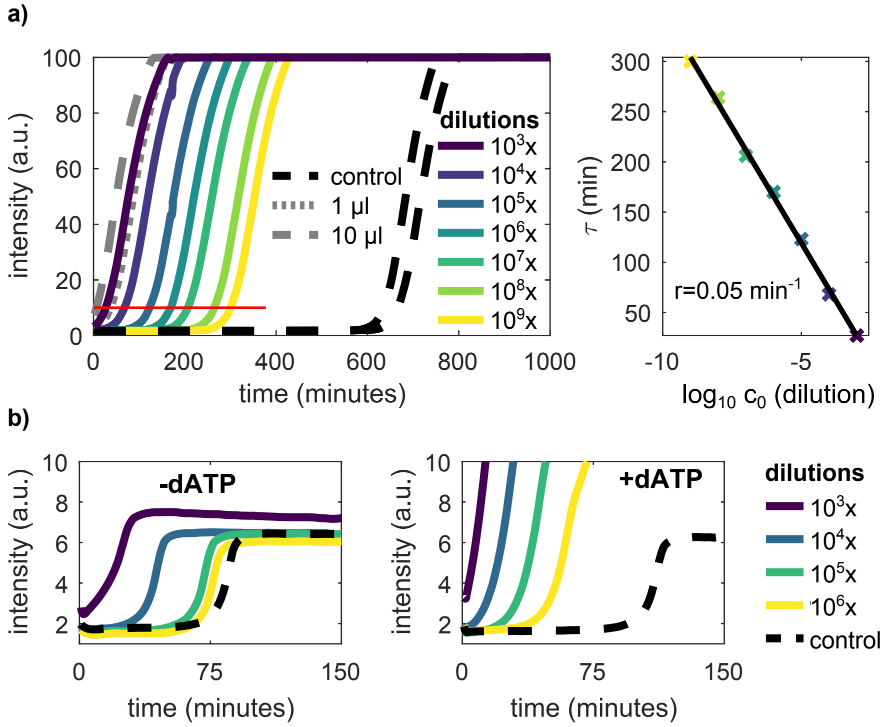


Figure S4: Amplification kinetics of parasite solutions diluted by different factors in the absence and in the presence of template strand. a) A mix containing all nucleotides and enzymes but no template was contaminated with different concentrations of parasite. A $10^9$-fold dilution can be easily distinguished from the non-contaminated control (black, dashed lines). For the grey curves, we infected pipette tips with parasite by either filling in 1 $\mu$l or 10 $\mu$l, emptying them and use them to mix a clean reaction mix. On the right panel we plotted the replication onset time $\tau$ *vs.* the logarithm of the dilution factor to determine the rate of parasite replication. b) In the presence of 50 nM $\mathbf{T}_1$ but no trigger $\mathbf{A}_1$, only very high concentrations of parasite ($\geq 10^4\times$ dilution) perturb rEXPAR (-dATP, left) while for standard EXPAR (+dATP, right) parasite concentrations as low as ($\geq 10^6\times$ dilution) make it impossible to distinguish EXPAR from parasite replication.

#### 6 Bistability

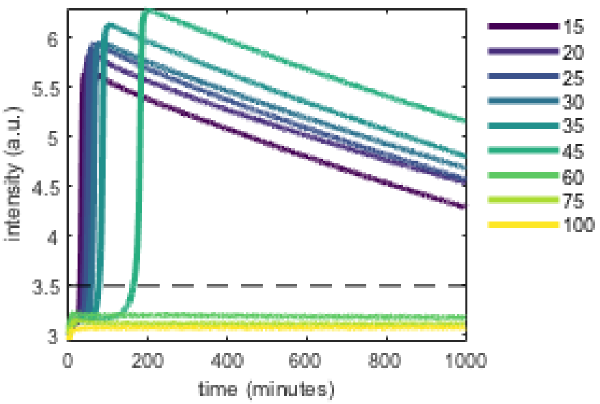


Figure 5: Full data corresponding to Figure 5 in the MT. EvaGreen fluorescence *vs.* time in the presence of $\mathbf{T}_1^u$ but in the absence of $\mathbf{A}_1$, for increasing concentrations of $\mathbf{R}_1$ in nM. Experiments performed in the absence of dATP. Conditions: 8 U/mL pol, 20 u/mL nick, 100 nM exo and 0.4 mM dNTPs. The signal decreases after amplification because the $\mathbf{T}_1^u$ template without protection against Nb.BssSI star activity was used.

### 7 Increasing nucleotide concentration slows down templated amplification

The removal of dATP from the buffer not only prevents parasite formation, but also speeds up the amplification reaction. Figure S6 shows the data for EXPAR experiments for different initial trigger concentrations. In Figure S6a and b on the left an EXPAR experiment is shown with and without dATP, respectively. A range of initial $\mathbf{A}_1$ concentration is replicated under otherwise identical conditions, with a trigger-free control (black dashed line). Comparing the graphs shows that the presence of dATP slows down the reaction. To quantify this effect, we plot the time $\tau$ it takes to reach a threshold intensity (red lines) versus the logarithm of the initial trigger concentration c0. From the slopes of the fitted lines we can determine the rates. We get $r = 0.90$ min$^{-1}$ in the presence of dATP and $r = 1.51$ min$^{-1}$ in its absence. This is a 1.66-fold increase in replication rate.

In the sequences we used, dATP is not incorporated during replication. Thus, one could argue that this effect is caused by the polymerase being hindered to perform its task by the unusable nucleotides being present in the buffer. However, we also observed a slowing down when the overall amount of nucleotides is reduced (Figure S7). This experiment was done at concentrations of 1 mM per nucleotide with dATP present or removed. In addition, time traces of reactions without dATP and a lower overall nucleotide concentration of 0.2 mM are shown (yellow dashed line). The right graph of the figure shows the initial 200 minutes in more detail. In agreement with the observations made above, the reaction is delayed in the presence of dATP. In addition, comparing the samples without dATP shows that lower dNTPs concentrations speed up the reaction.

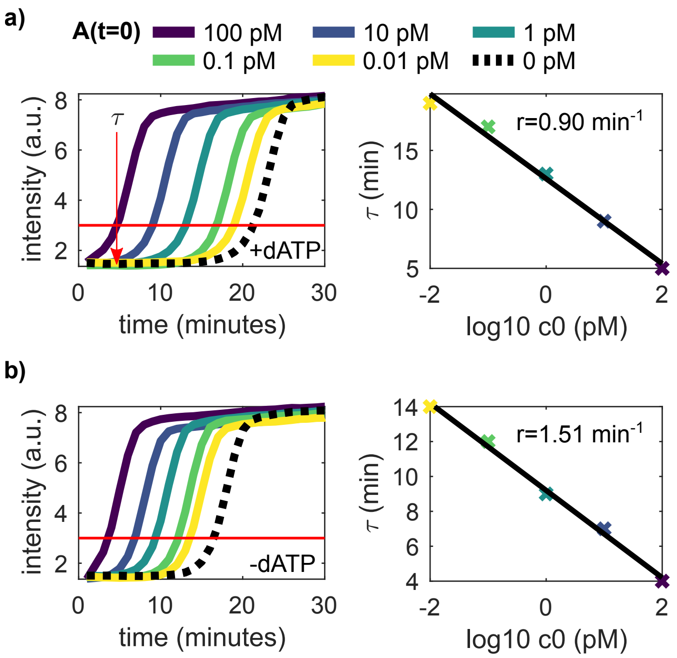


Figure S6: Templated amplification is faster in the absence of dATP. a) The left graph shows a range of initial $\mathbf{A}_1$ concentration undergoing an autocatalytic reaction in the presence of dATP. The red horizontal line gives a threshold intensity. The right graph shows the time $\tau$ it takes for each curve to reach the threshold. From this, the replication rate $r$ can be determined by a line fit. b) Shows the same experiment and analysis, but without dATP in the buffer.

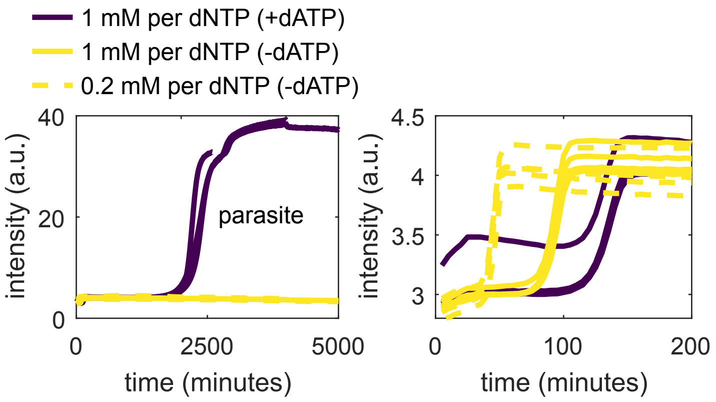


Figure S7: Effects of nucleotide composition and concentration on amplification kinetics. The left graph shows time traces for autocatalysis with 1 mM per nucleotide with and without dATP as well as autocatalysis with 0.2 mM per nucleotide without dATP. The samples were removed from the cycler at different times for PAGE analysis, thus only one time trace per condition stays in the cycler for the full length of the experiment. The right graph shows the first 200 minutes more detailed. The reaction speeds up when dATP is removed and when the overall nucleotide concentration is reduced. The reaction shown is untriggered and contains netrosin.

### 8 Amplification curves for EXPAR and rEXPAR in the presence of parasite contamination

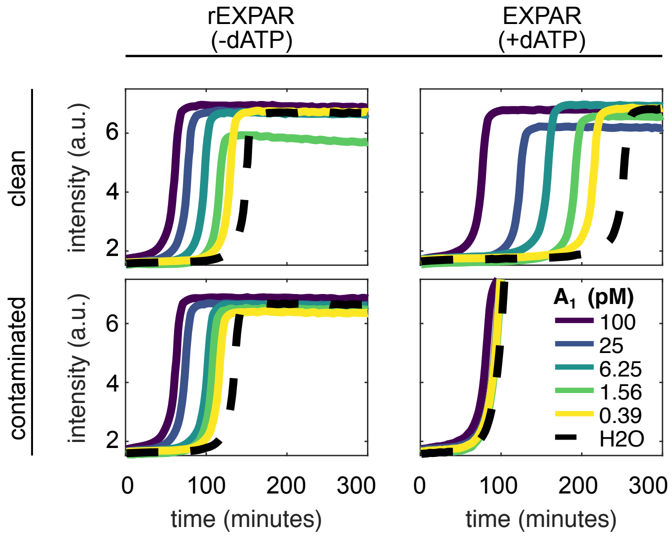


Figure S8: Amplification curves for EXPAR and rEXPAR in the presence of parasite contamination. These graphs show the raw data of Figure 7 of the main text. The threshold used for the analysis was 3.
